# How many days are needed to estimate accelerometry-assessed physical activity during pregnancy? Methodological analyses based on a cohort study using wrist-worn accelerometer

**DOI:** 10.1101/522722

**Authors:** Shana Ginar da Silva, Kelly R Evenson, Ulf Ekelund, Inácio Crochemore Mohsam da Silva, Marlos Rodrigues Domingues, Bruna Gonçalves Cordeiro da Silva, Márcio de Almeida Mendes, Glória Isabel Niño Cruz, Pedro Curi Hallal

## Abstract

**Background:** Objective methods to measure physical activity (PA) can lead to better cross-cultural comparisons, monitoring temporal PA trends, and measuring the effect of interventions. However, when applying this technology in field-work, the accelerometer data processing is prone to methodological issues. One of the most challenging issues relates to standardizing total wear time to provide reliable data across participants. It is generally accepted that at least 4 complete days of accelerometer wear represent a week for adults. It is not known if this same assumption holds true for pregnant women.

**Aim:** We assessed the optimal number of days needed to obtain reliable estimates of overall PA and moderate-to-vigorous physical activity (MVPA) during pregnancy using a raw triaxial wrist-worn accelerometer.

**Methods:** Cross-sectional analyses were carried out in the antenatal wave of the 2015 Pelotas (Brazil) Birth Cohort Study. Participants wore the wrist ActiGraph wGT3X-BT accelerometer for seven consecutive days. The daily average acceleration, which indicates overall PA, was measured as milli-*g* (m*g*), and time spent in MVPA (minutes/day) was analyzed in 5-minute bouts. ANOVA and Kruskal-Wallis tests were used to compare variability across days of the week. Bland-Altman plots and Spearman-Brown Prophecy Formula were applied to determine the reliability coefficient associated with one to seven days of measurement. Analyses were stratified by sociodemographic factors and nutritional status.

**Results:** Among 2,082 pregnant women who wore the accelerometer for seven complete days, overall and MVPA were lower on Sundays compared to other days of the week. Reliability of >=0.80 to evaluate overall PA was reached with at least three monitoring days, whereas six days were needed to estimate reliable measures of MVPA.

**Conclusions:** Our findings indicate that the usual approach obtaining one week of accelerometry in adults is also appropriate for pregnant women, particularly to obtain differences on weekend days and reliably estimate MVPA.

## Introduction

Objective methods to measure physical activity (PA), such as accelerometers, have become widely used over the years given the high degree of validity to assess patterns of PA in free-living conditions [1]. Accelerometry-based PA assessment can lead to better cross-cultural comparisons, monitoring temporal PA trends and measuring the effect of interventions [2]. However, when applying this technology in field-work, the accelerometer data processing is prone to methodological issues with important implications that can affect data quality [3,4]. One of the most challenging issues relates to standardizing total wear time to provide reliable data across participants [5–8].

Several studies have been carried out in children [9], young [10,11] and adult population [12,13] focused on the numbers of monitoring days necessary to represent habitual PA behavior. Results of these studies suggested a large variability in the number of days required to obtain reliable measures of PA ranging from 2 to 9 days. Also, the number of required days varies according to the intensity of physical activities, often grouped as sedentary behavior, and light, moderate, and vigorous intensity [12,13]. Other factors that can influence the monitoring time-frame are the type of accelerometer used and placement of the device (e.g., wrist, thigh, or waist) [6–8].

More recently, a growing interest in PA during pregnancy has emerged given the potential positive effects of PA on maternal-child health [14]. However, there are currently few studies which have used accelerometers to measure PA during pregnancy [15,16]. Moreover, research focus to determine a suitable monitoring time-frame to accurately measure PA behavior has been performed in young to middle-aged adults [10–13], and no data appear available among pregnant women. Therefore, the purpose of this study was to examine the optimal number of days needed to obtain reliable estimates of overall PA and MVPA during pregnancy using a raw triaxial wrist-worn accelerometer in a population-based study in southern Brazil. In addition, we aimed at measuring the variability in the means of PA across days of the week.

## Materials and Methods

### Design and participants

We conducted cross-sectional analyses based on the antenatal wave of the 2015 Pelotas (Brazil) Birth Cohort Study. Participants with an expected delivery date from January 1^st^ 2015 to 31^st^ December 2015 were eligible for the cohort and recruited from all health facilities offering antenatal care (public and private) in the city of Pelotas. Accelerometry data was collected between weeks 16 and 24 of gestation. Details regarding this study have been previously described elsewhere [17]. Ethical approval for this study was obtained from the Ethics Committee of the Physical Education School - Federal University of Pelotas, in accordance with official letter numbered 522/064, approved the study. All participants signed a written informed consent prior to participation.

### Measurements

The accelerometer used was the ActiGraph wGT3X-BT models (ActiGraph,Pensacola, FL, USA). These devices were lightweight (27 g) and compact (3.8 × 3.7 × 1.8 cm), allowing measurement of body movements over three orthogonal axes vertical (Y), horizontal right-left (X), and horizontal front-back axis (Z), within an acceleration dynamic range of ± 8g [18]. Participants wore the accelerometer on their non-dominant wrist (dorsally midway between the radial and ulnar styloid processes) during 24 hours for seven consecutive days. The accelerometer was programmed to collect raw acceleration at 60 Hz and three-dimensional raw data was expressed in gravitational equivalent units called milli-gravity (m*g*, where 1000m*g*= 1*g* = 9.81 m/s^2^).

### Data reduction

Devices were programmed and accelerometers’ data downloaded using ActiLife software, version 6.11.7. Accelerometer raw data analyses were performed in R-package GGIR [19]. The following parameters were used to consider valid data for the analyses: calibration error <0.02*g* and seven full days of measurement (total protocol). Euclidian Norm Minus One (ENMO) was used to summarize three-dimensional raw data (from axes x, y, and z) into a single-dimensional signal vector magnitude 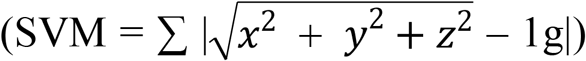 [19]. Data were further summarized when calculating the average values per 5-second epochs. The summary measures used were (a) overall PA (expressed in m*g*), based on the average SVM per day, (b) average time spent in MVPA per day with 5-minute bouts criterion (expressed in minutes). MVPA was defined as SVM records above 100m*g* [20,21], while bouts criterion was defined as consecutive periods in which participants spent at least 80% of time in activities with intensity equal or higher the MVPA threshold.

### Statistical analysis

Sample descriptions are presented in relative (%) and absolute frequencies (N). Overall PA was expressed in mean and standard deviation (SD), while MVPA was presented as a mean, SD, median, and interquartile range (25th and 75th percentiles). ANOVA and Kruskal-Wallis non-parametric test were used to compare whether PA varied significantly across days of the week. If an overall significant F level was shown, post-hoc tests (Bonferroni pairwise comparisons) were used to assess differences between weekdays (Monday to Sunday). The number of days required to reliably estimate habitual PA (overall PA and MVPA) was assessed using the Spearman-Brown formula. A modified version of the Spearman-Brown calculation determined the intraclass reliability coefficient associated with 1 to 7 days of measurement. The standard typically used for acceptable reliability is an ICC of >=0.80 [22]. We also assessed agreement based on the visual inspection of the Bland-Altman plots.

We stratified the analysis by maternal age (<20, 20-29, 30-39, ≥40), skin color (white, black, brown/yellow/indigenous), socioeconomic position (based on asset index [23] and later categorized into quintiles) paid job during pregnancy (yes/no), and pre-pregnancy body mass index (BMI) (calculated by dividing weight by height squared (kg/m^2^) with cutoffs were defined according to the World Health Organization [24]. All analyses were performed using Stata version 12.1 (StataCorp, College Station, TX, USA). Statistical significance was set at *α* < 0.05.

## Results

From 2,463 pregnant women with accelerometry data, 2,082 adhered to the research protocol and wore accelerometer for seven consecutive days. A high proportion of the sample was aged 20-29 (49.5%), white skin color (73.3%), did not have a paid job during pregnancy (50.1%), with normal pre-pregnancy BMI (48.8%) and belonged to the top quintile for socio-economic position (Table 1).

**Table 1.** Characteristics of participants that wore accelerometer for seven consecutive days. The 2015 Pelotas (Brazil) birth cohort study.

|  | n | % |
| --- | --- | --- |
| **Maternal age (years)** |  |  |
| <20 | 277 | 13.3 |
| 20-29 | 1,029 | 49.5 |
| 30-39 | 722 | 34.7 |
| ≥ 40 | 52 | 2.5 |
| **Skin color** |  |  |
| White | 1,523 | 73.3 |
| Black | 255 | 12.3 |
| Brown/yellow/indigenous | 299 | 14.4 |
| **SES (quintiles)** |  |  |
| Q1 (poorest) | 243 | 14.7 |
| Q2 | 327 | 19.8 |
| Q3 | 358 | 21.7 |
| Q4 | 359 | 21.7 |
| Q5 (wealthiest) | 366 | 22.1 |
| **Paid job during pregnancy** |  |  |
| No | 1,042 | 50.1 |
| Yes | 1,037 | 49.9 |
| **Pre-pregnancy BMI (Kg/m<sup>2</sup>)** |  |  |
| Underweight | 61 | 3.3 |
| Normal | 913 | 48.8 |
| Overweight | 536 | 28.7 |
| Obese | 360 | 19.3 |

SES: socioeconomic position; BMI: body mass index.

Mean overall PA (m*g*) and time spent in MVPA (minutes/day) was different across days of the week. Overall PA and time spent in MVPA was lower on Sunday (25.6 m*g* and 8.6 minutes/day, respectively) compared to all other days. Pregnant women were more physically active on weekdays (p<0.001) for both overall PA and MVPA (Table 2).

**Table 2.**
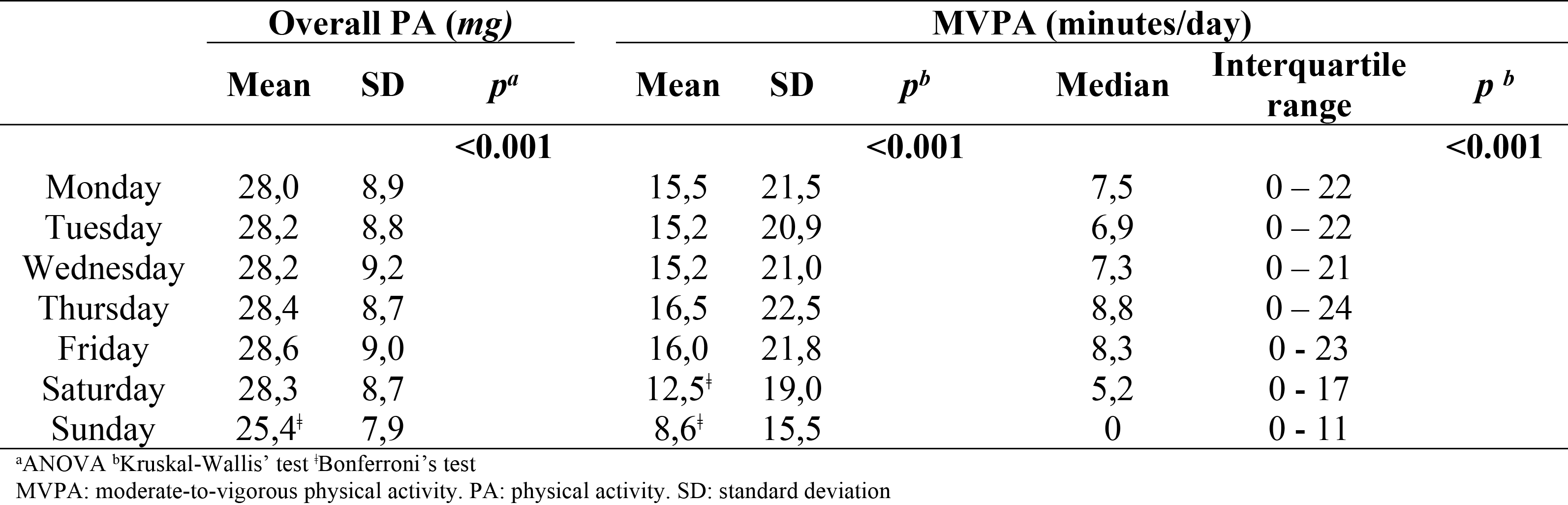
Daily duration (*mg* and minutes) of overall physical activity and moderate to vigorous physical activity

Estimates of the number of days needed to obtain reliable measures of habitual PA are presented in Figure 1.

**Figure 1.**
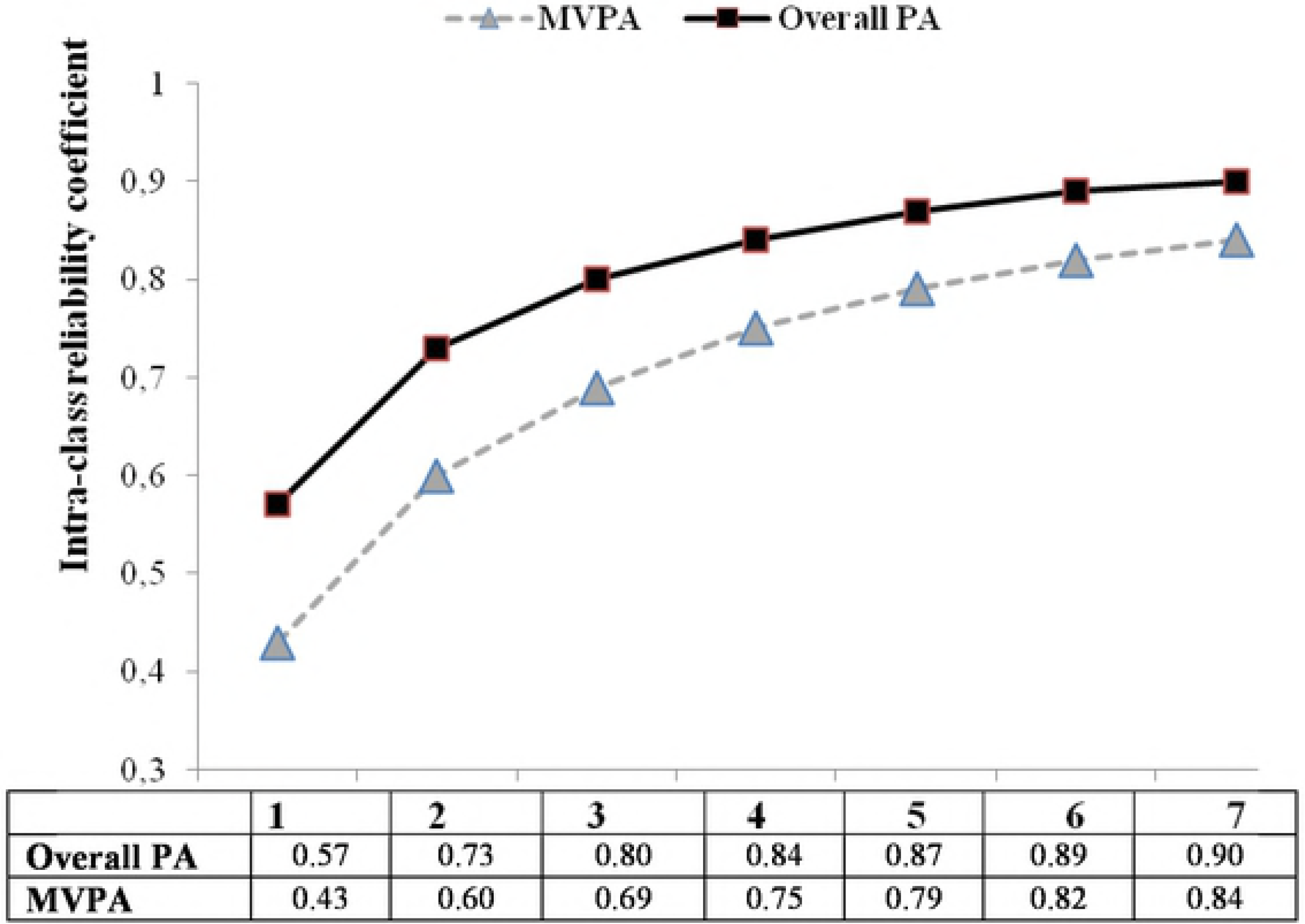
Reliability coefficient for number of days monitoring.

For overall PA, at least three days of the week was the minimum necessary to achieve a reliability of 0.80 whereas six monitoring days were needed to estimate reliable measures of MVPA. These results indicate that between 43–57%, 60–83%, 69–80%, 75–84%, 79-87%, 82-89% and 84-90% of the variance was accounted for using 1 to 7 days monitoring to represent habitual activity for overall PA and MVPA, respectively.

Table 3 presented the reliability coefficient associated with different number of monitored days stratified by age, nutritional status and sociodemographic groups. In terms of overall PA, a minimum of three days of monitoring show IRC (Intra-class reliability coefficient) values close to 0.80 for all groups of age, skin color, socioeconomic position, job characteristics and pre-pregnancy BMI. Reaching IRC values of around 0.80 requires a minimum of six days of use for MVPA, except for pregnant women younger than 20 years, who tend to have a more variable PA pattern and reach an IRC of 0.80 when monitored for at least seven days.

**Table 3.** Intraclass reliability correlation coefficient stratified by maternal age, SES and paid job during pregnancy in pregnant women belonging to the 2015 Pelotas (Brazil) Birth Cohort Study.

|  | Intraclass reliability coefficient by Spearman-Brown Prophecy Formula |  |  |  |  |  |  |  |  |  |  |  |  |  |
| --- | --- | --- | --- | --- | --- | --- | --- | --- | --- | --- | --- | --- | --- | --- |
|  | Overall PA |  |  |  |  |  |  | MVPA |  |  |  |  |  |  |
|  | 1 day | 2 days | 3 days | 4 days | 5 days | 6 days | 7 days | 1 day | 2 days | 3 days | 4 days | 5 days | 6 days | 7 days |
| **Maternal age (years)** |  |  |  |  |  |  |  |  |  |  |  |  |  |  |
| <20 | 0.52 | 0.69 | 0.77 | 0.81 | 0.85 | 0.87 | 0.88 | 0.29 | 0.45 | 0.55 | 0.62 | 0.67 | 0.71 | 0.74 |
| 20-29 | 0.57 | 0.72 | 0.80 | 0.84 | 0.87 | 0.89 | 0.90 | 0.45 | 0.62 | 0.71 | 0.76 | 0.80 | 0.83 | 0.85 |
| 30-39 | 0.61 | 0.76 | 0.82 | 0.86 | 0.89 | 0.90 | 0.92 | 0.45 | 0.62 | 0.71 | 0.77 | 0.80 | 0.83 | 0.85 |
| ≥ 40 | 0.44 | 0.61 | 0.70 | 0.76 | 0.80 | 0.82 | 0.85 | 0.41 | 0.59 | 0.68 | 0.74 | 0.78 | 0.81 | 0.83 |
| **Skin color** |  |  |  |  |  |  |  |  |  |  |  |  |  |  |
| White | 0.58 | 0.73 | 0.80 | 0.85 | 0.87 | 0.89 | 0.91 | 0.43 | 0.61 | 0.70 | 0.75 | 0.79 | 0.82 | 0.84 |
| Black | 0.57 | 0.73 | 0.80 | 0.84 | 0.87 | 0.89 | 0.90 | 0.43 | 0.60 | 0.69 | 0.75 | 0.79 | 0.82 | 0.84 |
| Brown/Yellow/Indigenous | 0.55 | 0.71 | 0.78 | 0.83 | 0.86 | 0.88 | 0.89 | 0.37 | 0.54 | 0.64 | 0.70 | 0.75 | 0.78 | 0.81 |
| **SES (quintiles)** |  |  |  |  |  |  |  |  |  |  |  |  |  |  |
| Q1(poorest) | 0.59 | 0.74 | 0.81 | 0.85 | 0.88 | 0.90 | 0.91 | 0.45 | 0.62 | 0.71 | 0.76 | 0.80 | 0.83 | 0.85 |
| Q2 | 0.53 | 0.69 | 0.77 | 0.82 | 0.85 | 0.87 | 0.89 | 0.39 | 0.56 | 0.66 | 0.72 | 0.76 | 0.79 | 0.82 |
| Q3 | 0.60 | 0.75 | 0.82 | 0.86 | 0.88 | 0.90 | 0.91 | 0.39 | 0.56 | 0.65 | 0.72 | 0.76 | 0.79 | 0.82 |
| Q4 | 0.55 | 0.71 | 0.79 | 0.83 | 0.86 | 0.88 | 0.89 | 0.36 | 0.53 | 0.63 | 0.69 | 0.74 | 0.77 | 0.80 |
| Q5 (wealthiest) | 0.56 | 0.72 | 0.80 | 0.84 | 0.87 | 0.89 | 0.90 | 0.41 | 0.58 | 0.68 | 0.74 | 0.78 | 0.81 | 0.83 |
| **Paid job during pregnancy** |  |  |  |  |  |  |  |  |  |  |  |  |  |  |
| No | 0.58 | 0.73 | 0.80 | 0.84 | 0.87 | 0.89 | 0.90 | 0.39 | 0.56 | 0.65 | 0.72 | 0.76 | 0.79 | 0.82 |
| Yes | 0.57 | 0.72 | 0.80 | 0.84 | 0.87 | 0.89 | 0.90 | 0.48 | 0.65 | 0.73 | 0.78 | 0.82 | 0.85 | 0.86 |
| **Pre-pregnancy BMI (Kg/m<sup>2</sup>)** |  |  |  |  |  |  |  |  |  |  |  |  |  |  |
| Underweight | 0.57 | 0.73 | 0.80 | 0.84 | 0.87 | 0.89 | 0.90 | 0.42 | 0.59 | 0.69 | 0.74 | 0.78 | 0.81 | 0.84 |
| Normal | 0.58 | 0.73 | 0.80 | 0.85 | 0.87 | 0.89 | 0.91 | 0.45 | 0.63 | 0.71 | 0.77 | 0.81 | 0.83 | 0.85 |
| Overweight | 0.58 | 0.71 | 0.81 | 0.85 | 0.88 | 0.89 | 0.91 | 0.42 | 0.59 | 0.69 | 0.75 | 0.79 | 0.81 | 0.84 |
| Obese | 0.55 | 0.71 | 0.78 | 0.83 | 0.86 | 0.88 | 0.89 | 0.42 | 0.59 | 0.69 | 0.74 | 0.78 | 0.81 | 0.84 |

SES: socioeconomic position; MVPA: moderate-to-vigorous physical activity; PA: physical activity; BMI: body mass index.

Bland-Altman plots indicated on average differences between number of days near zero, narrow limits of agreement and a homogeneous variability across the days of monitoring, for both overall PA and MVPA. Thus, more days of monitoring produce lower variability between measurement days (1 to 6) and the standard seven-day protocol for both MVPA and overall PA (Figure 2 and 3).

**Figure 2.**
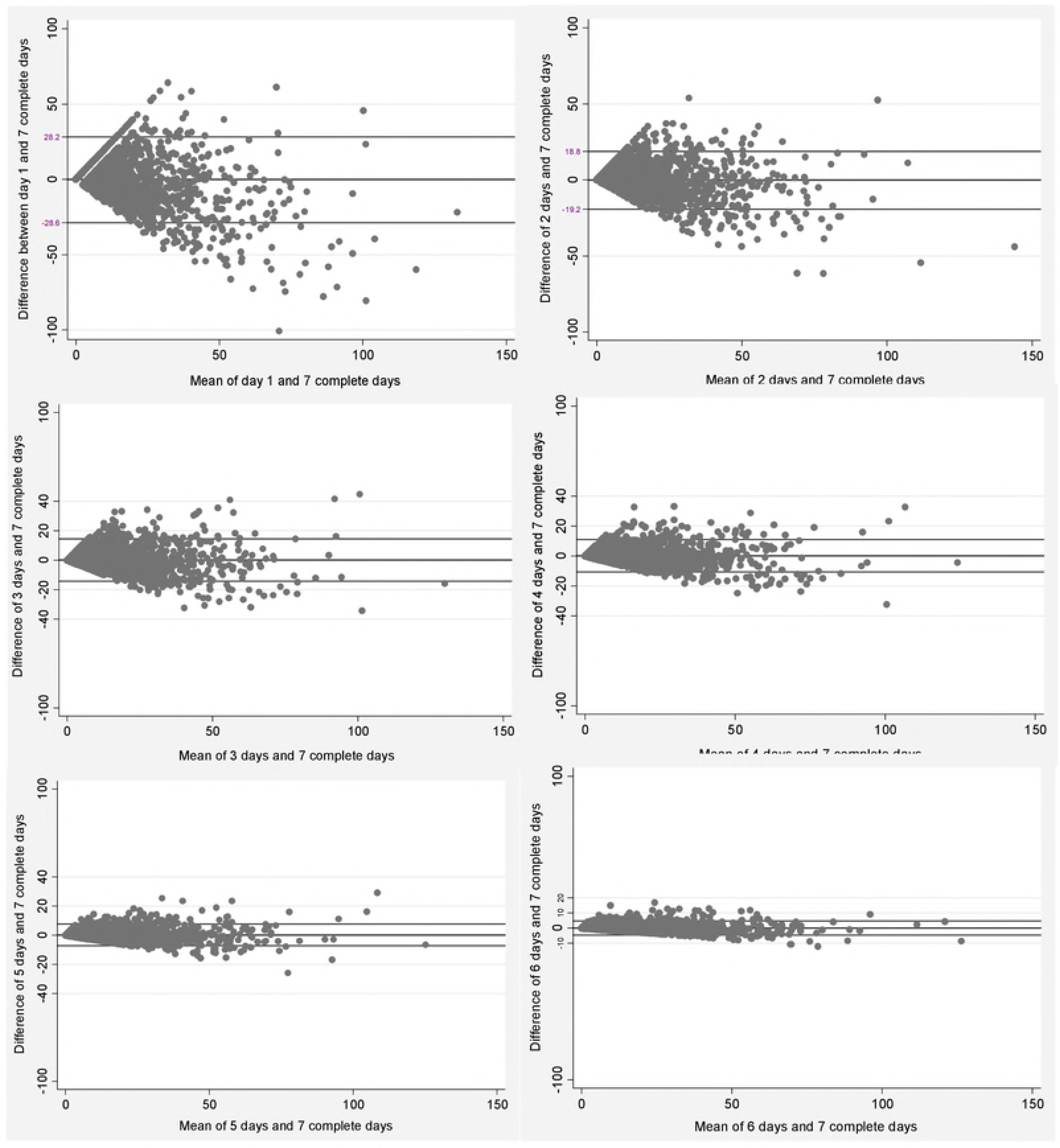
Bland-Altman plots of the comparison between the means of measurement days (1 to 6) and the standard of seven complete days of measurement for moderate-to-vigorous physical activity (MVPA).

**Figure 3.**
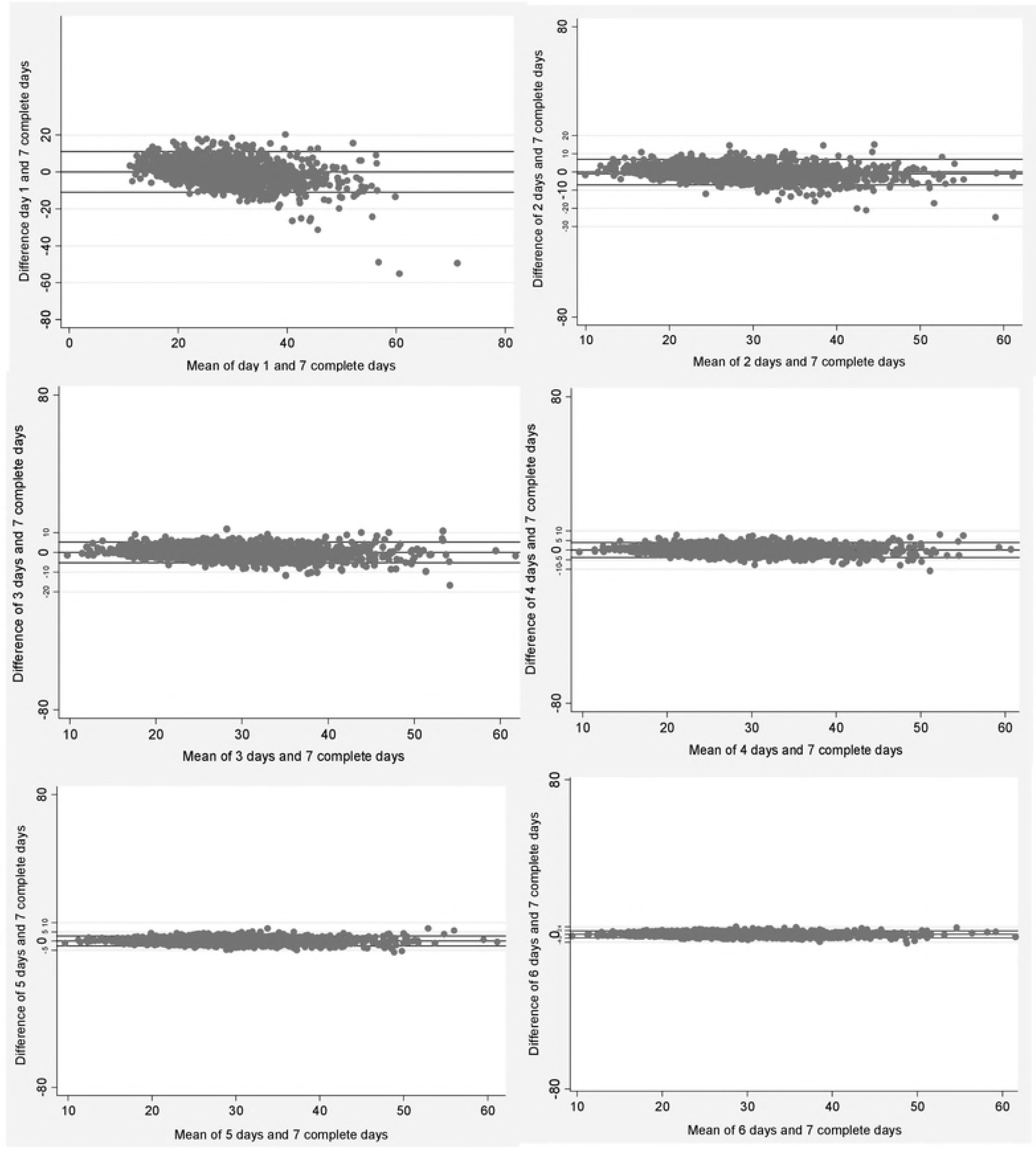
Bland-Altman plots of the comparison between the means of measurement days (1 to 6) and the standard of seven complete days of measurement for overall physical activity.

As expected, higher mean differences were found between one day and seven complete days for both MVPA (mean difference: 0.36; 95%CI: -0.31-1.02) and overall PA (mean difference: 0.09; 95%CI: -0.15; 0.33). On the other hand, lower mean differences were identified between six days of measurement and the standard protocol in the two intensities investigated, MVPA (mean difference: -0.11; 95%CI: -0.21; -0.01) and overall PA (mean difference: -0.03; 95%CI: -0.06; 0.01), respectively.

## Discussion

This study determined the number of monitoring days needed to obtain reliable estimates of overall PA and MVPA in pregnant women using wrist-worn accelerometers in a population-based study in southern Brazil. Our findings showed that at least six monitoring days of the week should be considered to achieve a reliability of 0.80 to accurately predict both overall PA and MVPA levels. Variability in the means of overall PA and MVPA across the days of the week was also observed and clear differences were noted, with the lowest means of overall PA and MVPA on Sunday. This finding indicates that weekend days cannot be ignored in the design and analysis of PA studies. Considered together, these findings support the usual approach of asking adults to wear an accelerometer for one week.

To the best of our knowledge, this is the first study to date to investigate the number of days needed to obtain reliable estimates of overall PA and MVPA during pregnancy in a representative population sample using raw triaxial wrist accelerometry. Thus, our observations are not directly comparable with previous observations. However, it seems well established in the literature that the number of days needed to obtain reliable estimates of habitual activity varies according to the intensity of physical activities measured. A study conducted by Dillon and cols [12], using wrist-worn GENEActiv accelerometers investigated an acceptable reliability measure of weekly habitual PA in middle-aged Irish adults. They also found that the monitoring frame duration for reliable estimates varied across intensity categories. Results ranged from 2 days when evaluating combined MVPA to 6 days for specifically vigorous activities. Matthews et al. [25] using the Computer Science Applications (CSA) accelerometer on the hip in healthy adults determined that 3–4 days monitoring were required to accurately measure MVPA. Similar results were reported by Hart et al. [13] in a study with older adults using waist-worn accelerometers. Contrary to these findings, we observed that six monitoring days are necessary to produce reliable measures of MVPA among pregnant women. Pregnancy is a complex period that involves many physical and psychological changes including morphological adjustments for an ideal environment for fetal development, changes in mood, anxiety, fatigue/energy, etc [26]. These factors may contribute to a larger variability in MVPA measurements throughout the week in pregnant women compared to other populations.

Previously published data have indicated a wide variability in the number of days needed to estimate a reliable measure of PA in different population groups. Several aspects may explain the inconsistency between prior results such as the heterogeneity in the type of accelerometer adopted, number of accelerometers used in the studies (single or multiple body position), available budget for data collection and placement of the device (hip/waist or wrist). Another question that may influence the differences is the statistical techniques applied to obtain stable mean estimates of PA. These discrepancies in the methods of each study emphasize the need to establish an appropriate monitoring frame to reliably capture habitual physical behavior for each population, accelerometer, PA intensity and body position of wearing the devices [27].

Patterns of PA during pregnancy are influenced by demographic, economic, environmental and behavioral characteristics [15, 27]. Considering the possible influence of these aspects on the number of days required to represent weekly habitual PA, analyses were stratified by sociodemographic factors and nutritional status. Similar results were found for all groups except for pregnant women younger than 20 years, who needed more than 7 days of monitoring to achieve reliable measures of MVPA.

The valid and reliable activity monitor, 24-hour study protocol, large sample size, high-rate response, wrist-worn accelerometer and statistical techniques employed are strengths of our study. The Spearman-Brown prophecy formula has been used in most studies investigating appropriate monitoring frames [12]. However, some limitations should be noted. In our study, accelerometers were used for seven complete and consecutive days. Monitoring for longer periods, such as a month, season or a year, would be alternatives to obtain greater representativeness of habitual PA behavior given that many studies have reported seasonal and monthly variations in PA [29, 30]. However, a longer collection time would probably result in lower compliance and bring logistic issues during collection (such as battery replacement and data downloading). Also, our results showed that measuring six consecutive days we could reliably estimate overall PA and MVPA in this group of pregnant women.

In addition, our findings are not advocating for future studies among pregnant using only three (to estimate overall PA) or six monitoring days (to estimate MVPA). Our results suggest that a seven day protocol may be optimal when assessing habitual PA in pregnant women. If a short time of assessment is applied, there will be no room for non-wear time, which might lead for a limited number of valid data. Furthermore, it is important to note that our analyses presented the minimum necessary for a reliable estimate of habitual PA objectively measured, which might be specific for our research context. These set of analyses are highly recommended for each single study in order to robustly define their inclusion criteria in terms of minimum of valid days.

## Conclusion

Our results indicate that at least three days of monitoring are required to reliably capture overall PA and six days monitoring when considering MVPA. Due to the substantially lower PA levels during Sundays, we recommend a seven day protocol throughout a full week when assessing habitual PA in pregnant women. These findings may have implications for future study designs and data reduction strategies among accelerometer-assessed physical activity studies of pregnant women.

## Acknowledgments

The authors would like to thank the Wellcome Trust, the Brazilian National Research Council (CNPq), and the Coordination for the Improvement of Higher Education Personnel (CAPES) for providing financial support to this study. da Silva SG would like to thank you CAPES and CNPq (n. 439505/2016-0) for the scholarship and financial support.

